# Delaying quantitative resistance to pesticides and antibiotics

**DOI:** 10.1101/2022.08.14.503896

**Authors:** Nate B. Hardy

## Abstract

How can we best vary the application of pesticides and antibiotics to delay resistance evolution? Previous theoretical studies have focused on qualitative resistance traits, and have mostly assumed that resistance alleles are already present in a population. But many real resistance traits are quantitative, and the evolution of resistant genotypes in the field may depend on *de novo* mutation and recombination. Here, I use an individual-based, forward-time, quantitative-genetic simulation model to investigate the evolution of quantitative resistance. I evaluate the performance of four application strategies for delaying resistance evolution, to wit, the (1) sequential, (2) mosaic, (3) periodic, and (4) combined strategies. I find that which strategy is best depends on initial efficacy. When at the onset, xenobiotics completely prevent reproduction in treated demes, a combined strategy is best. On the other hand, when populations are partially resistant, the combined strategy is inferior to mosaic and periodic strategies, especially when resistance alleles are antagonistically pleiotropic. Thus, the optimal application strategy for managing against the rise of quantitative resistance depends on pleiotropy and whether or not partial resistance is already present in a population. This result appears robust to variation in pest reproductive mode and migration rate, direct fitness costs for resistant phenotypes, and the extend of refugial habitats.

## Introduction

Pest and pathogen populations evolve resistance to pesticides and antiobiotics. For example, in the last 80 years, in more than 500 insect species, almost 8,000 cases of insecticide resistance have been reported to more than 300 compounds (Denholm, Devine, & Williamson, 2002; Mota-Sanchez & Wise, 2022). Likewise, among more than 1,700 bacteria species, tens of thousands of putative resistance alleles have been characterized, conferring resistance to more than 200 antibiotics (Liu & Pop, 2009). In principle, one way to manage resistance evolution would be to keep developing new xenobiotics. But that might not be sustainable. It currently takes about ten years and $1 billion to develop a new antibiotic, and it has been several decades since new insecticide or herbicide modes of action were discovered (Gaines, Busi, & Küpper, 2021; REX, 2013). The fact is that we have been losing effective chemical controls of pest populations faster than we have replaced them (Gaines et al., 2021). This is already taking a toll on public health and prosperity (Friedman, Temkin, & Carmeli, 2016). It could become worse. For antibiotics, by one estimate, before 2050, antimicrobial resistance will have killed 2.4 million people in Europe, North America and Australia, and cost the world economy $3.5 billion in losses per year (OECD, 2018). If sustainable development of novel xenobiotics is not feasible, then what we can do to conserve the ones we have?

To be clear, in the field, one of the strongest predictors of xenobiotic resistance is application intensity (Evans et al., 2015; Hicks et al., 2018); effective xenobiotic conservation will undoubtedly entail using them more sparingly and shifting to more integrated pest management approaches (Mascanzoni et al., 2018; Massa et al., 2013). But that leaves the question of how, when we do use xenobiotics, we should vary their application over time and space so as to maximize their durability, and thus the time a pest population is under control. That is our focus here. Supposing for simplicity’s sake that we have just two pesticides, we have four main conservation strategies to consider: (1) *Sequential application (a*.*k*.*a*., *responsive alternation)*. Use one pesticide everywhere until resistance evolves, then switch to the second. This is the default practice in agriculture. The upside is that when the second pesticide is deployed, the pest population is naive to it. Of course, the downside is that selection for resistance to each pesticide is unremitting. (2) *Periodic application (a*.*k*.*a*., *rotation)*. Like sequential application, this is a way of varying treatments over time but not space. But rather than start with one pesticide and wait until resistance evolves before changing to the second, we switch back and forth between pesticides at some arbitrary frequency. If genotypes that are resistant to one pesticide are susceptible to the other, such rotations should impede the rise of alleles conferring resistance to either. On the other hand, across a few generations, periodic application presents both adaptive targets to a pest population, and this might boost selection for cross-resistance genotypes. (3) *Mosaic application*. Vary applications in space but not time. This strategy is used more often in medicine than agriculture (Reluga, 2005). If pests move between treatment demes, this could establish a selective regime similar to that of periodic application; with an individual as the frame of reference, migration between treatment areas equates to temporal variation in selection. But varying pesticides over space rather than time could change epidemiological dynamics and the population genetic structure of pest populations in a way that affects adaptation (Plantegenest, Le May, & Fabre, 2007). (4) *Combined (a*.*k*.*a*., *mixed or pyramided) application*. The logic behind this strategy is quite different from the previous three. Instead of trying to delay resistance evolution by adding heterogeneity to the selective environment, we use both pesticides in concert to consistently apply the strongest selection possible.

So, which is best? The evidence for pesticide resistance was reviewed last by The Resistance to Xenobiotics Consortium (REX, 2013). Since then, several additional studies have been published, mostly considering how to conserve the efficacy of *R-*genes in cultivated plant varieties (e.g., Djian-Caporalino et al., 2014; Finckh et al., 2000; Nilusmas et al., 2020; Rimbaud, Papaïx, Rey, Barrett, & Thrall, 2018; Sudo, Takahashi, Andow, Suzuki, & Yamanaka, 2018; reviewed by Rimbaud et al., 2021), along with some additional studies of herbicide and antibiotic resistance (Busi, Powles, Beckie, & Renton, 2019; Evans et al., 2015; Tepekule, Uecker, Derungs, Frenoy, & Bonhoeffer, 2017). Empirically, we do not have much to go on; experimental comparisons of the evolutionary ramifications of application strategies are no easy task, and almost all of empirical work has been fraught by problems with experimental design (no replication, no randomization, etc.) and has examined pest populations with standing high-frequency resistance alleles (as noted by REX, 2013; but see Lagator, Vogwill, Colegrave, & Neve, 2012 for an exception). Likewise, comparative analyses of the evolution of resistance in the field (as recommended by Comont & Neve, 2021) is all but precluded by poor records of the history of local and regional pesticide use (but see Hicks et al., 2018 for example of this kind of analysis). More progress has been made on the theory side; indeed, with the above-mentioned recent work on *R*-gene conservation and herbicide resistance, there have now been more than 40 published studies that directly compare two or more xenobiotic application or *R*-gene deployment strategies. (Note that this is a subset of more expansive work on the evolution of resistance evolution, most of which does not directly compare alternative conservation strategies (REX, 2010)). For the most part, this theory indicates that the combined strategy should be best and the sequential strategy should be the worst, with the rankings of the periodic and mosaic strategies depending on system-specific features. Taken at face value, managers should always use a combined application strategy. Of course, that might not be economical; indeed, theoretical evaluations of resistance management strategies have seldom considered economics (REX, 2010; but see (Ndeffo Mbah, Forster, Wesseler, & Gilligan, 2010). Moreover, to my knowledge, all published theoretical comparisons of have assumed that resistance traits are qualitative, that is, that resistance is due to large-effect alleles at one or two loci affecting target-site insensitivity. This may be a useful approximation of some systems, but in many cases key resistance traits are quantitative, that is, resistance is due to small-effect alleles at many genetic loci affecting metabolism, behavior or xenobiotic penetrance (Comont et al., 2022; Crossley, Chen, Groves, & Schoville, 2017; Delourme et al., 2006; Hδllinger, Pennings, & Hermisson, 2019; Mohammad, Osborne, & Freckleton, 2022). Likewise, most studies have assumed that resistant genotypes are already present in pests populations, but in the field resistance may depend on *de novo* mutation and recombination.

Hence, there is a gap in our understanding of how to delay resistance, specifically, for cases in which resistance is quantitative. Here, to begin to fill in this gap, I develop an individual-based, stochastic, forward-time quantitative-genetic simulation model. I then use the model to compare the performance of the sequential, mosaic, periodic, and combined strategies, across a variety of demographic scenarios, pest life histories, and quantitative genetic architectures for resistance.

## Methods

Simulations were performed with SLiM 3 (Haller & Messer, 2019). In brief, SLiM simulates the forward-time evolution of a population of individual genomes which can reproduce, mutate, recombine, and die. Population-level properties and processes, such as populations size and genetic drift, emerge from individual-level properties and processes, such as viability and mating. It consists of a set of set of C++ libraries wrapped by a higher-level programming language called Eidos that abstracts away many of the details of the simulated life cycle and evolutionary process, and allows researchers to focus on the specification of genetic architectures, demographic scenarios, populations structures, and selective regimes. The Eidos code for the model described next are available via GitHub (S1; https://github.com/n8-rd/resistance).

### Initialization

Each simulation starts with a population of ***N***=6,000 hermaphroditic individuals, spread evenly across 30 demes. Suppose that these individuals are herbivorous insects, and that demes are farms. The starting population is monomorphic; each individual has the same diploid 100kb, one-chromosome genome, with a quantitative genotype value of zero for each of two resistance traits (described below). After initializing the population, we iterate over generations of the life cycle (Fig. 1).

**Figure 1.**
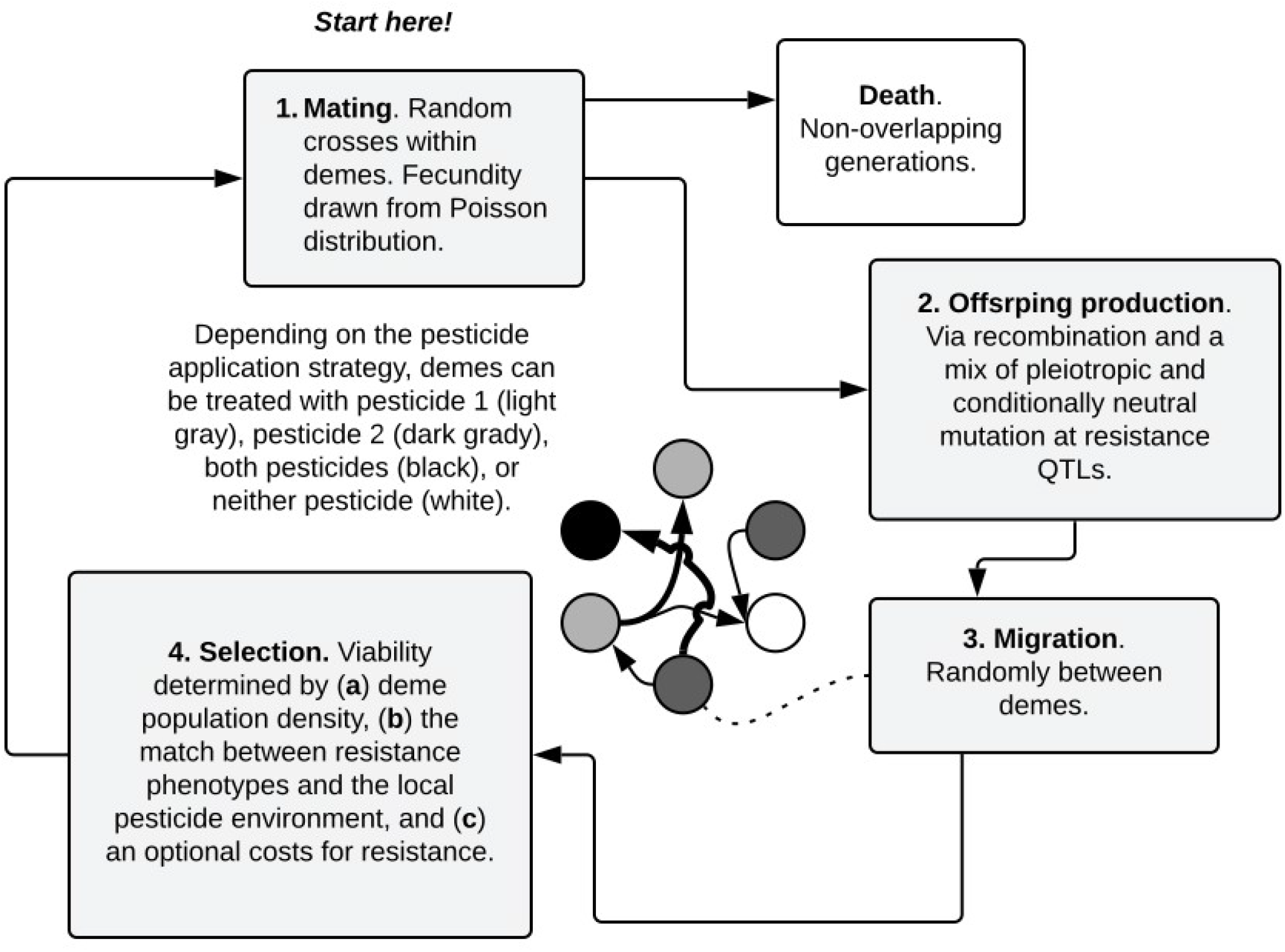
The simulated pest life cycle. Each generation consists of (1) a round of random, within-deme mating, with fecundity determined by a draw from a random Poisson distribution with ***λ***=1, (2) offspring production entailing recombination of parental genomes, and quantitative mutation, with allele effects drawn from random normal distributions, (3) random migration among 30 demes, and (4) viability selection, in which and individual’s probability of surviving to the mating stage is determined by (a) deme density, (b) the match between their resistance phenotypes and the local pesticide application strategy, and (c) direct viability costs for resistance phenotypes. To be clear, in the model, pesticide application takes place after migration but before mating.

#### 1. Mating

The first step is mating, which is monogamous. Each surviving individual randomly selects a mate from their deme. Individual fecundity is determined by a draw from a Poisson distribution, with a mean and variance of one. Generations do not overlap, that is, the population is semelparous; after offspring production, parents die.

#### 2. Offspring production

After mating comes offspring production, which entails recombination of parental genomes as well as mutation. To reiterate, the finer details of how mutation and recombination happen are taken care of by SliM. The recombination rate is set at 1e-8 events per site per generation, which maps to approximately ten crossing-over events per genome per generation – about what we would expect for a real herbivorous insect species with a five-chromosome genome (Wilfert, Gadau, & Schmid-Hempel, 2007). There are three types of mutation: (1) conditionally neutral mutations that affect resistance to pesticide 1 but not pesticide 2, (2) conditionally neutral mutations that affect resistance to pesticide 2 but not pesticide 1, and (3) pleiotropic mutations that affect both resistance phenotypes. The conditionally neutral mutation rate is ten times that of pleiotropic mutation, 1e-6 versus 1e-7 per site, per individual, per generation; in nature this disparity might be yet more extreme (Gompert et al., 2015), but a 10:1 ratio at least roughly approximates the preponderance of conditional neutrality, while leaving the door open for pleiotropy. When a conditionally neutral mutation occurs an allele effect is drawn from a univariate normal distribution with a mean of zero, and a variance of 0.5. When a pleiotropic mutation occurs, a two-dimensional allele effect vector is drawn from a bivariate normal distribution with means of zero, variances of 0.5, and symmetrical covariances which varied across simulations, taking one of the values −0.45, 0, or 0.45. These allele effects are part of what determines viability, that is, the probability that an individual will survive long enough to reproduce. Details are provided below.

#### 3. Migration

Before viability selection each new individual has a chance of migrating to another deme, given by parameter ***m***. As a robustness check, I examined several values for ***m*** (Table 1), but as this had little effect on the relative performance of application strategies, I focus on models in which ***m*** was set to 0.1. Migration among demes is completely random, that is, there is no explicit spatial structure, and migrants do not evaluate destinations, so all migration targets are equally likely. Although this assumption is a simplifying expediency, it may well approximate some real systems, for example where population mixing is enhanced by the trade of infected nursery stock, or unsanitary farming equipment.

**Table 1.**
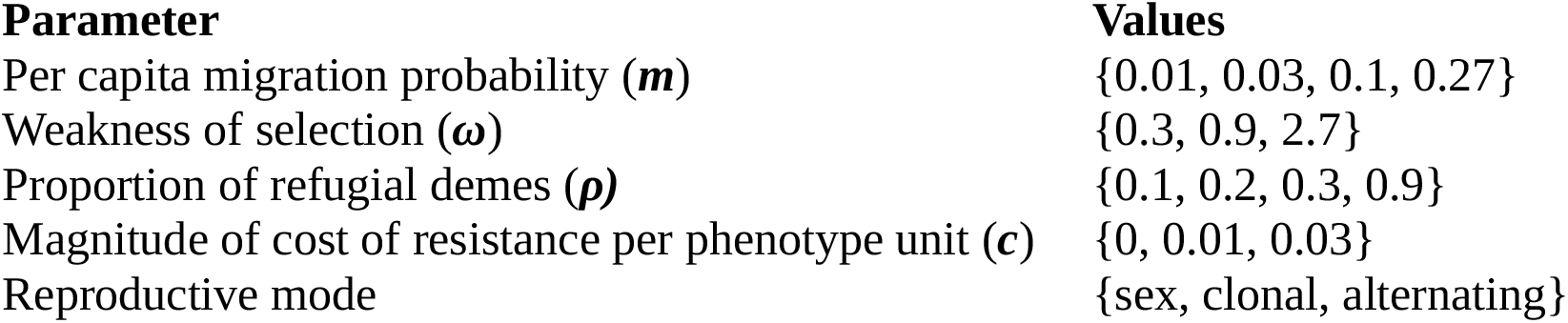
Model parameters that varied across simulations.

| Parameter | Values |
| --- | --- |
| Per capita migration probability ($m$) | {0.01, 0.03, 0.1, 0.27} |
| Weakness of selection ($\omega$) | {0.3, 0.9, 2.7} |
| Proportion of refugial demes ($\rho$) | {0.1, 0.2, 0.3, 0.9} |
| Magnitude of cost of resistance per phenotype unit ($c$) | {0, 0.01, 0.03} |
| Reproductive mode | {sex, clonal, alternating} |

#### 4. Selection

The next step in the life cycle is selection. In broad strokes, an individual’s viability is determined by three factors: (1) deme population density, (2) phenotype-environment matching, specifically, the match between an individual’s resistance phenotypes and the local pesticide application regime, and (3) an optional cost for resistance phenotypes. Next, I describe each of these factors in more detail.

#### 4a. Density dependence

I start each simulation with a short burn-in period in which the population accumulates genetic diversity at resistance-affecting sites before we apply any pesticides. Concretely, for the first 49 generations, all alleles are neutral, and individual viability is determined solely by local population density. To be clear, in the model, deme population size is not fixed, but rather emerges via the balance of individual mortality and fecundity. Each deme has a nominal carrying capacity, ***K***_***i***_, of ***N***/30=200 individuals. Individual viability is scaled by the ratio of ***K***_***i***_ to the current local subpopulation size, ***K***_***i***_/***N***_***i***_. I allow this scaling factor to vary from zero to 1.5, but note that an individual’s viability cannot be greater than one. Thus, high population density within a deme reduces the viability of all genotypes, but low population density only boosts the viability of maladapted genotypes. A biological justification for this fitness boost at low density would be escape from apparent competition, which is thought to be key limiting factor on herbivorous insect populations (Bird, Kaczvinsky, Wilson, & Hardy, 2019). Note that by fixing ***K***_***i***_ values, I assume an aseasonal, perennial cropping system. Relaxing this aseasonality assumption – and periodically setting the ***K***_***i***_ of treated demes to zero – has little effect on model dynamics (results not shown); for simplicity’s sake, I focus on the aseasonal model.

#### 4b. Phenotype-environment matching

At generation 50, pesticide application commences. From that point on, viability is determined in part by an individual’s quantitative resistance phenotypes. Each individual has two resistance traits: resistance to pesticide 1, and resistance to pesticide 2. The first resistance genotype value is equal to the sum of allele effects at sites that are pleiotropic or that affect the pesticide 1 resistance phenotype but not pesticide 2 resistance. Likewise, the second resistance genotype value equals the sum of corresponding pleiotropic and conditionally neutral allele effects. Resistance phenotypes are determined by adding to each of these genotype values an environmental effect, which in each generation is drawn from a normal distribution with mean zero and variance of 0.2. In mathematical terms,

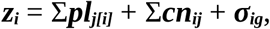

where ***z***_***i***_ is an individual’s resistance phenotype ***i, pl***_***j[i]***_ is the quantitative effect on phenotype ***i*** of an allele at a pleiotropic locus ***j, cn***_***ij***_ is the effect on phenotype ***i*** of an allele at a conditionally neutral locus ***j***, and ***σ***_***ig***_ is the environmental effect on phenotype ***z***_***i***_, in generation ***g***.

Viability is determined by the matches between an individual’s resistance phenotypes and the local resistance optima in their deme (Fig 2). Which optima apply depends on the local pesticide application strategy. Individuals in a deme may be exposed to (1) pesticide 1, (2) pesticide 2, (3) both pesticides, or (3) no pesticides, that is, they may occur in a refuge. As for migration rates, I explored a range of values for the proportion of demes which are refuges, ***ρ***, but focus on models in which the proportion is 0.2 (Table 1).

**Figure 2.**
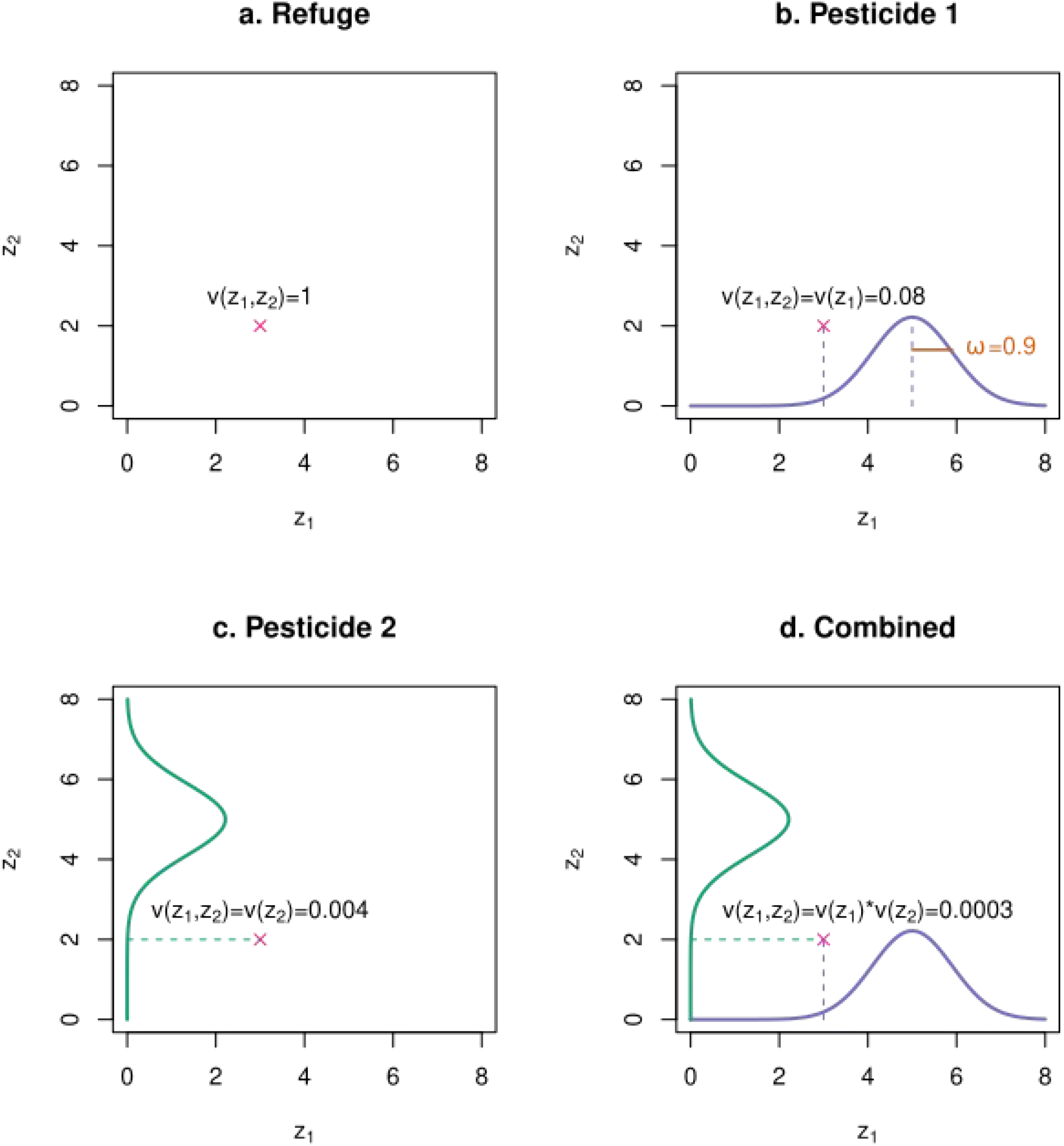
Schematic of the mapping of resistance phenotypes to viability, desregarding effects from deme density and possible direct costs for resistance. The **x** denotes a hypothetical individual’s two-dimensional resistance phenotype, with the pesticide 1 resistance phenotype, ***z***_***1***_, given on the x-axis and the pesticide 2 resistance phenotype, ***z***_***2***_, on the y-axis. **a**. In a refuge, there is no selection on resistance and therefore, all phenotypes have a viability, ***v***(***z***_***1***_,***z***_***2***_), of one. **b, c**. In demes exposed to only one pesticide, and individual’s corresponding resistance phenotype determines its viability via a Gaussian fitness function; viability is maximized (and equals one) when a resistance phenotype perfectly matches the optimum value (here, set to five). The variance of the Gaussian distribution, ***ω***, gives the weakness of selection. **d**. In demes exposed to a combination of both pesticides, an individual’s overall viability is equal to product of its single-component viabilities, ***v***(***z***_***1***_,***z***_***2***_)=***v***(***z***_***1***_)\****v***(***z***_***2***_).

The mapping of phenotypes to viability effects is by a Gaussian fitness function:

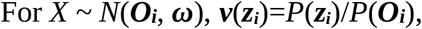

where ***z***_***i***_ is an individual’s phenotype value for resistance against pesticide ***i, v***(***z***_***i***_) is the viability of an individual with ***z***_***i***_, ***O***_***i***_ is the pesticide-specific optimal value, ***ω*** is the variance, that is, the weakness of selection (smaller variances equate to steeper selection gradients), and division by *P*(***O***_***i***_) sets the viability of a perfect match to one. In non-mathematical terms, individuals with more optimal resistance phenotype have higher viability, that is, a better chance of surviving long enough to mate. In each generation, the optimum resistance phenotype against pesticide 1, ***O***_***1***_, is 5 + ***d***, and the optimum resistance phenotype against pesticide 2, ***O***_***2***_, is also 5 + ***d***, where ***d*** is a random deviate from a zero-centered Gaussian distribution with variance of 0.2. Again, translating to non-mathematical terms, resistance optima are distant from the population’s starting resistance genotype values (ten standard deviations of the allele affect distribution) and vary a little bit from one generation to the next. In the main model – which has no direct fitness costs for resistance phenotypes – in demes where individuals are exposed to only one pesticide, allele effects governing the other resistance trait are neutral. Resistance trait-specific viabilities combine multiplicatively, that is ***v***(***z***_***1***_,***z***_***2***_) = ***v***(***z***_***1***_) * ***v***(***z***_***2***_). (Fig. 2 includes an example calculation.)

#### 4c. Direct costs for resistance phenotypes

The last viability component is a cost for resistance, ***C***. This cost is proportional to the summed resistance phenotype values, that is, ***C*** = ***c***(***z***_***1***_+***z***_***2***_), where ***c*** is the cost scaling factor, with values given in Table 1. ***C*** is subtracted from an individual’s viability, determined already by deme density and phenotype-environment matching and subject to the constraint that a viability cannot be less than zero.

Putting it all together, accounting for density-dependence, phenotype-environment matching, and direct costs for resistance, the viability of individual ***i*** in deme ***j*** is ***v***_***i***_ = ***v***(***z***_***1***_,***z***_***2***_)(***K***_***j***_/***N***_***j***_) - ***c***(***z***_***1***_+***z***_***2***_); subject to 0 < ***K***_***j***_/***N***_***j***_ < 1.5 and 0 < ***v***_***i***_ < 1.

After viability selection, the life cycle begins again, with mating between surviving individuals.

What we want to know is which application strategy maximizes the duration of effective pest control. To that end, I define effective control – somewhat arbitrarily – as ending with the first appearance of a pest genotype value within 50% of the optimum value for either pesticide. For each simulation, I logged how many generations it takes for such a genotype to arise.

As previously mentioned, as robustness checks I examined the effects of varying pleiotropic allele effect covariances, migration rates (***m***), the proportion of refuges (***ρ***), and the magnitude of direct viability costs for resistance phenotypes (***c***). I also examine the impact of switching to clonal or alternating sexual-clonal modes of reproduction, with one round of sexual production for every tend rounds of clonal reproduction (Table 1). I ran twenty replicated simulations for each configuration of model parameters.

## Results and Discussion

### Relative timing of resistance evolution

Let us start with a close look at a base model in which reproduction is sexual, the proportion of refugial demes is 0.2, the migration rate is 0.1, and there are no direct fitness costs for resistance phenotypes. (Fig. 3). The sequential strategy is always a poor option. Which strategy is best depends on the strength of selection, that is, the initial efficacy of a control strategy.

**Figure 3.**
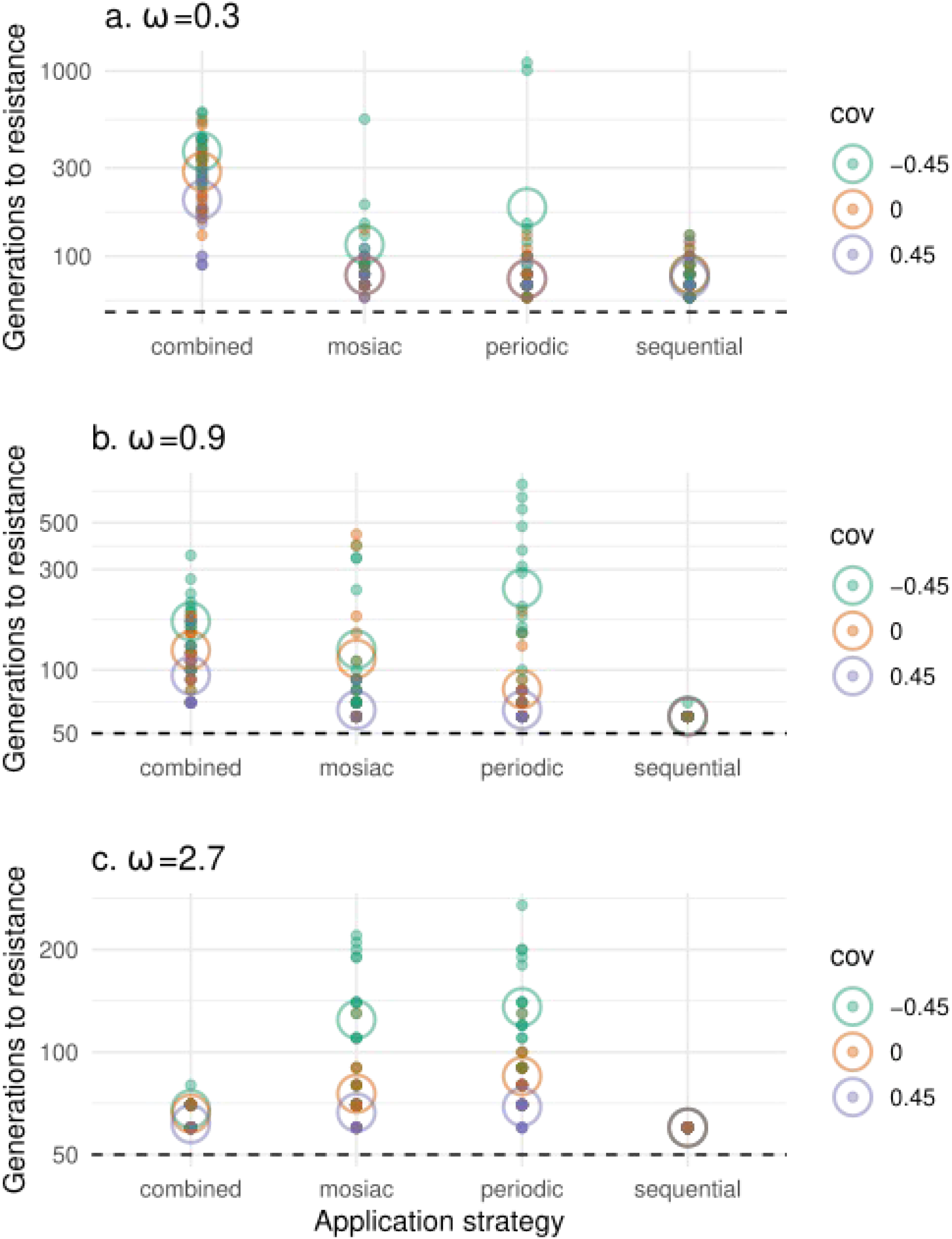
Main model comparison of application strategies. The position of each point on the y-axis shows, for a specific simulation, the number of generations until the emergence of first resistance. Points are color coded by pleiotropic covariance (cov): teal for −0.45, orange for 0, and lavender for 0.45. The y-axis is in on a logarithmic scale. Open circles show the mean values for a particular pleiotropic covariance. The dashed horizontal line shows the onset of pesticide application, at generation 50. Each plot shows the result of 20 simulations under each combination of application strategy and pleiotropic covariance. The parameter, ***ω***, gives the weakness of selection. *Interpretation*: As we go from strong to weak selection, and especially with strong negative pleiotropic covariances, we reverse the relative efficacy of the combined versus the mosaic and periodic application strategies.

When selection is strong, ***ω*** = 0.3 (Fig. 3a), regardless of pleiotropic covariances, the combined application strategy is the top performer (at ***α***=0.001, according to a linear model in which the response variable is generations until resistance and the predictor variables are application strategy, pleiotropic covariance, and the interaction between strategy and pleiotropic covariance). In comparison to the mosaic and combined strategies, it increases the mean time until first resistance by a factor of ∼5 (328 generations with the combined strategy versus 59 for each the mosaic and periodic strategies). Note that the superiority of the combined strategy in this case is congruent with what is typically found in theoretical studies of qualitative resistance evolution (REX, 2013).

But when selection is weak, ***ω*** = 2.7 (Fig. 3c), and therefore the pest population starts off partially resistant, the combined strategy is inferior to the mosaic and periodic strategies (Fig. 3b,c). Just how inferior depends on the covariances of pleiotropic alleles. To clarify, with strongly positive covariances, cross-resistance alleles prevail. When that is the case, performance differences between strategies are relatively small. With strongly negative covariances, pleiotropy is mostly antagonistic, and increased resistance to one pesticide tends to come at the cost of reduced resistance to the other. In this case, I find the the mosaic and periodic strategies do much better than the combined strategy; this is especially true of the periodic strategy, which in comparison to the combined strategy increases the mean time until first resistance by a fact or ∼7 (82 generations for the periodic strategy versus 11 for the combined).

Under medium-strength selection, ***ω*** = 0.9 (Fig. 3b), I find something in between the two extreme cases. When antagonistic pleiotropy prevails, the combined, mosaic and periodic strategies are about equal on average, although the variance of resistance times is tighter under the combined strategy. On the other hand, when pleiotropic covariances are zero or positive, the combined strategy outperforms the rest.

So, as we go from strong to weak selection, we see a reversal in the relative efficacy of the combined strategy versus the mosaic and periodic strategies. When a pest population lacks partial resistance, the combined strategy is best. But when a pest population has partial resistance, the mosaic and periodic strategies are better options.

### The dose debate

Although I had not anticipated it, this finding has ramifications for the so-called “dose debate,” that is, the question of how changing pesticide dose will affect the rate and nature of resistance evolution (reviewed by Neve, Busi, Renton, & Vila-Aiub, 2014). In brief, reducing pesticide dose reduces the kill rate, and thus the strength of hard selection, while also shrinking the minimum allele effect size likely to have an appreciable impact on fitness. This is expected to decrease the rate at which qualitative resistance evolves via large-effect alleles, while at the same time increase the rate of quantitative resistance evolution via combinations of small-effect alleles (Renton, Diggle, Manalil, & Powles, 2011).

The model here adds nuance. To wit, Figure 4 shows that the effect of dose – reflected by ***ω*** – depends on application strategy. With the combined and sequential strategies, weaker selection speeds up quantitative resistance evolution. In polynomial regressions of time-to-resistance against ***ω***, under the combined or sequential strategies, the first order ***ω*** effect is negative, the second order effect is positive, and both terms are significant (***α***=0.0001); thus, there is a convex but monotonically decreasing relationship between ***ω*** and the time until resistance. In contrast, under the mosaic and periodic strategy, ***ω*** has little effect on resistance evolution times, although under the periodic strategy there is a weak hump-shaped relationship, with resistance evolution taking a little longer on average when selection is medium-strength (second order ***ω*** effect negative and significant at ***α***=0.1). The upshot is that the mosaic and periodic approaches are more robust against variation in pesticide dose – an insight that may be especially useful for managers seeking to reduce agricultural inputs.

**Figure 4.**
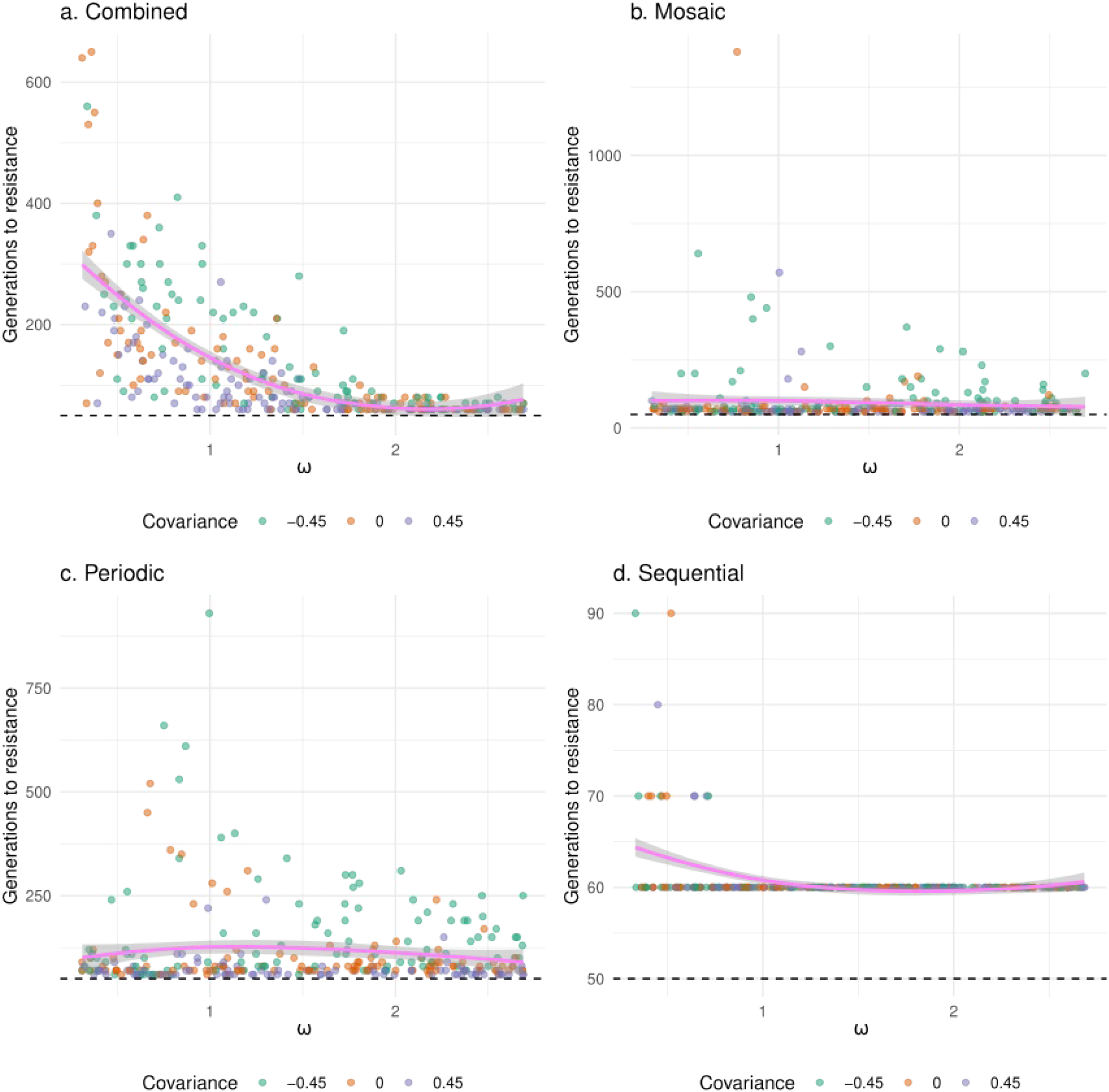
Effect of pesticide dose on resistance evolution. For these plots, in each simulation, the value of ***ω*** – which can be seen as a reflection of pesticide dose – was drawn from a random uniform distribution ranging from 0.3 to 2.7. As in Fig. 3, the color of each point corresponds to the pleiotropic covariance structure in effect for a simulation. The magenta line shows the trend across all pleiotropic covariances. *Interpretation*: For the combined and sequential strategies, the time it takes for a simulation pest population to evolve resistance is a monotonically increasing and concave function of dose. (Parameter ***ω*** is the weakness of selection; larger values correspond to weaker selection, and hence lower doses.) In contrast, under the mosaic and periodic strategies, the evolution of resistance is little affected by pesticide dose.

### Absolute timing of resistance evolution

Now that we have considered the relative performance of strategies, let me say few words about the absolute timing of resistance evolution. To be clear, the times reported here are a somewhat arbitrary function of the rates of mutation and recombination, the distribution of mutational effects, and the distance of optimal resistance phenotype values from the phenotypes of initialized pest populations. I did not attempt to match model parameter values to any particular pathosystem, but I did choose combinations of parameter values caused dynamics that at least roughly approximate what happens in the field, with resistance evolution taking from a few tens to a few hundreds of generations (Mota-Sanchez & Wise, 2022). Some readers may find the upper end of this spectrum too high. Here, are a couple of justifications.

First, empirical records of resistance times are “right censored” (Clark, Bradburn, Love, & Altman, 2003), that is, much of the history of resistance evolution is yet to unfold, and many of the longest waiting times have yet to be recorded. Second, the model shows what might happen if an application strategy is implemented consistently across a system, but in the field, application strategies tend to vary from one location to the next, and such heterogeneity can accelerate resistance evolution; previous theoretical studies of qualitative resistance have shown that the durability of a combined application strategy can be much diminished when combined treatment areas are embedded in a mosaic of single-component treatment areas, or when combined treatments follow single-component treatments (Lof, de Vallavieille-Pope, & van der Werf, 2017; Sapoukhina, Durel, & Le Cam, 2009). So, notwithstanding the right-censoring of the empirical resistance data, some of the longer resistance times simulated here can be taken as evidence of the control were are losing for lack of more coordinated regional-level pest management programs (Evans et al., 2018).

### Robustness checks

#### Reproductive mode

When resistance is qualitative and polymorphic, some classical population genetics theory predicts that resistance evolution should happen more rapidly in clonal than sexual pest populations (Crowder & Carrière, 2009). But here – where resistance is quantitative and pest populations are initially monomorphic for a genotype quite a distance from the optimum – I find the opposite; resistance evolution is much slower to evolve in clonal (and alternating clonal/sexual) populations (Suppl. Fig. 2). That being said, the relative performance of resistance delaying strategies is little affected by reproductive mode; the combined strategy is best when selection is strong, and the mosaic and periodic strategies are better options when it is weak.

#### *Migration rate (*m)

With sexual reproduction, and ***ρ***=0.2, the biggest effect is that increasing migration tends to increase the rate of resistance evolution across all strategies, especially when selection was weak (Suppl. Fig. 3). Changing migration rates does not affect the dependence of the optimal application strategy on the strength of selection; with strong selection, combined application is best, whereas with weak selection, the mosaic and periodic strategies are better options.

#### *Proportion of refugial habitats (*ρ)

With sexual reproduction, and ***m***=0.1, increasing the the proportion of refugial demes tends to delay resistance across strategies and selective strengths (Suppl. Fig. 4), as per previous theoretical and empirical work (Andrews, Li, & Lovenheim, 2016; Gould, 1998; Jin et al., 2015; Tabashnik, Gassmann, Crowder, & Carrière, 2008). The ***ω***-dependent relative performance of strategies is similar to that of the base model, except that under medium-strength or weak selection, with values for ***ρ*** above or below that used in the base model (0.2), the mosaic strategy outperforms the periodic strategy (Suppl. Fig. 4). Why the relative performance of mosaic and strategies would have such a complex mapping to ***ρ*** I cannot say, although previous work has shown that the efficacy of periodic strategies depends on the effective amplitude and period of selection (Reluga, 2005), while the efficacy of mosaic strategies depends on the relative carrying capacities of treated and untreated areas (Djidjou-Demasse, Moury, & Fabre, 2017; Lenormand & Raymond, 1998).

#### *Costs for resistance* (C)

As has been previously shown for polygenic, gene-for-gene, qualitative resistance (Bourget, Chaumont, & Sapoukhina, 2013), increasing the cost for resistance tends to slow down resistance evolution across pesticide application strategies, but has little effect on the patterns of relative performance we found in the base model (Suppl. Fig. 5). One exception is that with weak selection and a steep fitness cost for resistance, the periodic strategy is far superior to the mosaic strategy. Another exception is that with medium-strength selection and a steep fitness cost for resistance, the combined strategy is superior across pleiotropic covariances. Part of the logic of the combined strategy is that it pushes the two-dimensional optimum resistance phenotype far enough away from the pest population mean that most resistance-affecting mutations have a negligible effect on viability. It seems that at the medium-strength level of selection examined here, with non-combined strategies, initial pest genotype values for resistance are close enough to the optimum that the viability gains of resistance alleles are sufficient to offset their direct fitness costs. That seems not to be the case with the combined strategy.

In sum, the relative performance of resistance-delaying strategies is fairly robust to variation in reproductive mode, migrations rate, the proportion of refugial demes, and direct fitness costs for resistance phenotypes. Nevertheless, there is some evidence of complex and consequential interactions among those parameters.

### Caveats and contextualizations

As in any modeling exercise, I sought to strike a productive balance between specific realism and generalizability, which in this case was closer to the latter end of that spectrum. Therefore, further work – comparing strategies to delay quantitative resistance evolution with more system-specific parameterizations – would be useful. Likewise, the model makes many assumptions, for example, about population structure and the mapping of resistance phenotypes to fitness, and further work relaxing these assumptions, would also be useful.

This is not the first demonstration that the efficacy of a strategy to delay the evolution of xenobiotic resistance depends the population genetic context. A similar dependency was found for qualitative resistance under two-locus “gene-for-gene” architectures: a combined strategy outperforms periodic and mosaic strategies only when populations initially lack resistance-affecting alleles (Lof et al., 2017; Rimbaud, Papaïx, Luke, et al., 2018). And epidemiological compartment models of antibiotic resistance show that the efficacy of a combined treatment strategy can depend on the *de novo* rate at which double resistance emerged in patients (Tepekule et al., 2017). Likewise, this is not the first study to demonstrate that the performance of a strategy for delaying resistance can depend on genetic architecture (Bourget, Chaumont, Durel, & Sapoukhina, 2015). Thus, the model here can be seen as corroborating the results of several previous models. Nevertheless, this work departs from what has come before by explicitly comparing of strategies for delaying resistance evolution when resistance is quantitative. In that regard it addresses a critical gap in our understanding of resistance evolution. It may also serve as a springboard for investigations of the epigenetic evolution of resistance (Gressel, 2009; Mohammad et al., 2022).

## Conclusions

What works best for delaying the evolution of resistance to xenobiotics when resistance phenotypes are quantitative is not always the same as for when resistance is qualitative. Moreover, when resistance is quantitative, the best way of conserving effective pest control depends on whether or not a pest population is already partially resistant. Concretely, without partial resistance, the combined application strategy is the best option for keeping it that way. But with partial resistance, the mosaic and periodic application strategies are better options. When it comes to conserving what is in our xenobiotic toolkit, the genetic architectures of resistance phenotypes matter, as do prevailing population genetic conditions.

## Data archiving

There are no data to share, but the simulation model codes are provided as Supplementary material (S1).

## Supplementary Information

**S1**. Code for the simulation model.

**Figure S2**. Sensitivity of application strategies to variation in reproductive mode. The weakness of selection is given by parameter ***ω***. The position of each point on the y-axis shows, for a specific simulation, the number of generations until the emergence of first resistance, on a logarithmic scale. Points are color coded by pleiotropic covariance (cov): teal for −0.45, orange for 0, and lavender for 0.45. Open circles show the mean values for a particular pleiotropic covariance. The dashed horizontal line shows the onset of pesticide application, at generation 50.

**Figure S3**. Sensitivity of application strategies to variation in, ***m***, the migration rate. The weakness of selection is given by parameter ***ω***. The position of each point on the y-axis shows, for a specific simulation, the number of generations until the emergence of first resistance, on a logarithmic scale. Points are color coded by pleiotropic covariance (cov): teal for −0.45, orange for 0, and lavender for 0.45. Open circles show the mean values for a particular pleiotropic covariance. The dashed horizontal line shows the onset of pesticide application, at generation 50.

**Figure S4**. Sensitivity of application strategies to variation in, ***ρ***, the proportion of refugial demes. The weakness of selection is given by parameter ***ω***. The position of each point on the y-axis shows, for a specific simulation, the number of generations until the emergence of first resistance, on a logarithmic scale. Points are color coded by pleiotropic covariance (cov): teal for −0.45, orange for 0, and lavender for 0.45. Open circles show the mean values for a particular pleiotropic covariance. The dashed horizontal line shows the onset of pesticide application, at generation 50.

**Figure S5**. Sensitivity of application strategies to variation in, ***c***, the cost of resistance phenotypes. The weakness of selection is given by parameter ***ω***. The position of each point on the y-axis shows, for a specific simulation, the number of generations until the emergence of first resistance, on a logarithmic scale. Points are color coded by pleiotropic covariance (cov): teal for −0.45, orange for 0, and lavender for 0.45. Open circles show the mean values for a particular pleiotropic covariance. The dashed horizontal line shows the onset of pesticide application, at generation 50.

