## Supplementary figures and images for "Delaying quantitative resistance to pesticides and antibiotics"

### S2

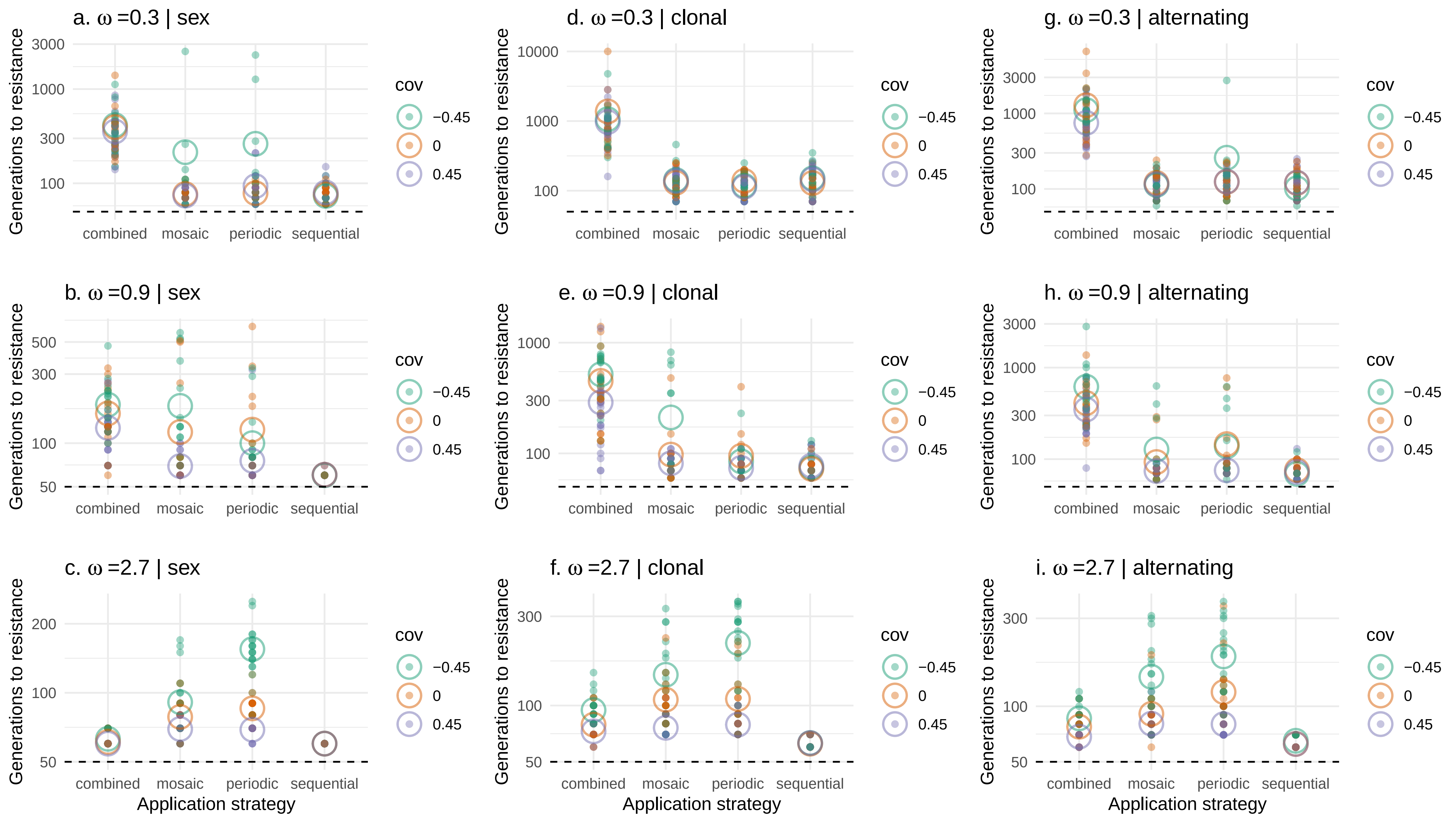

### S3

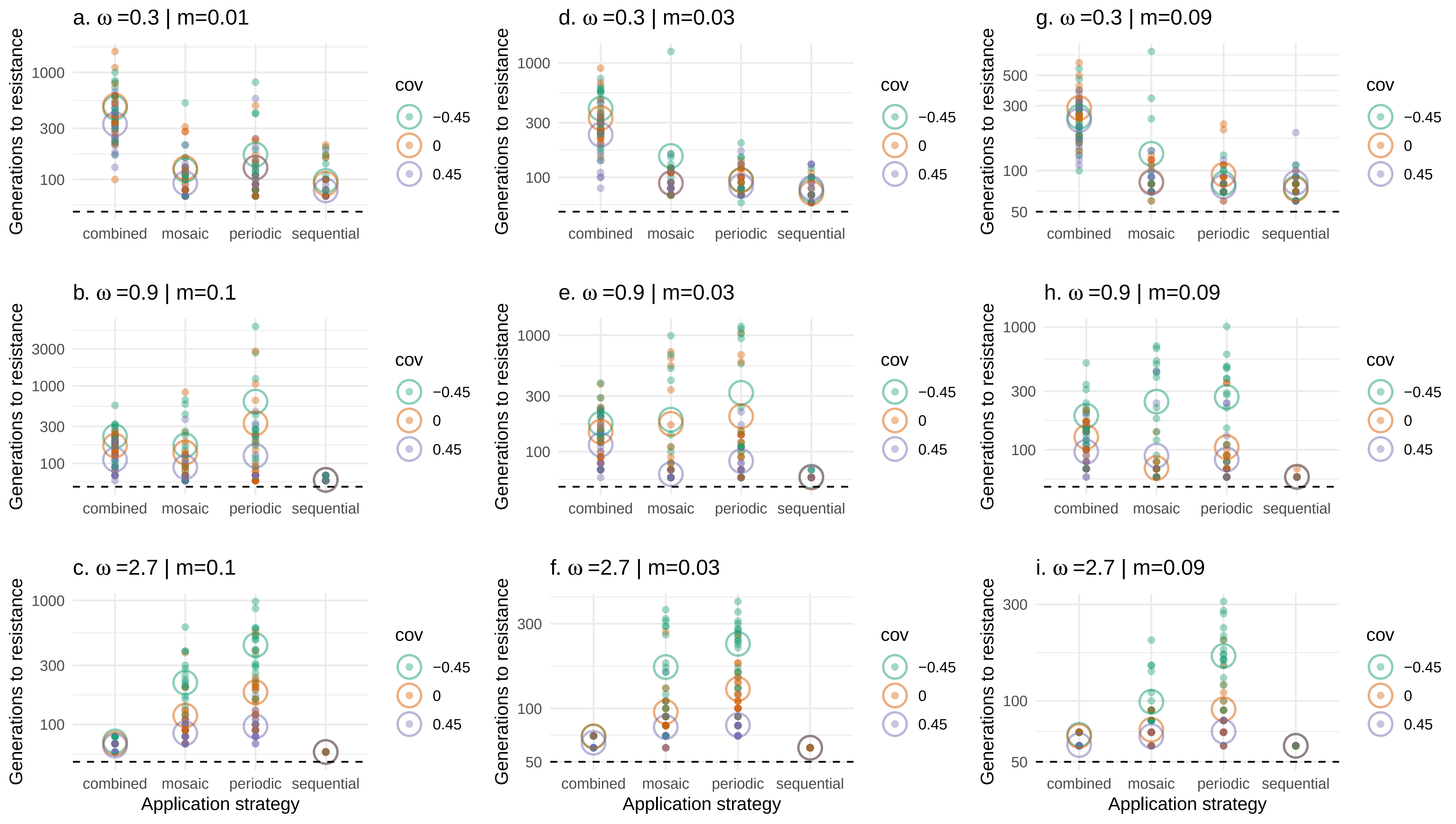

### S4

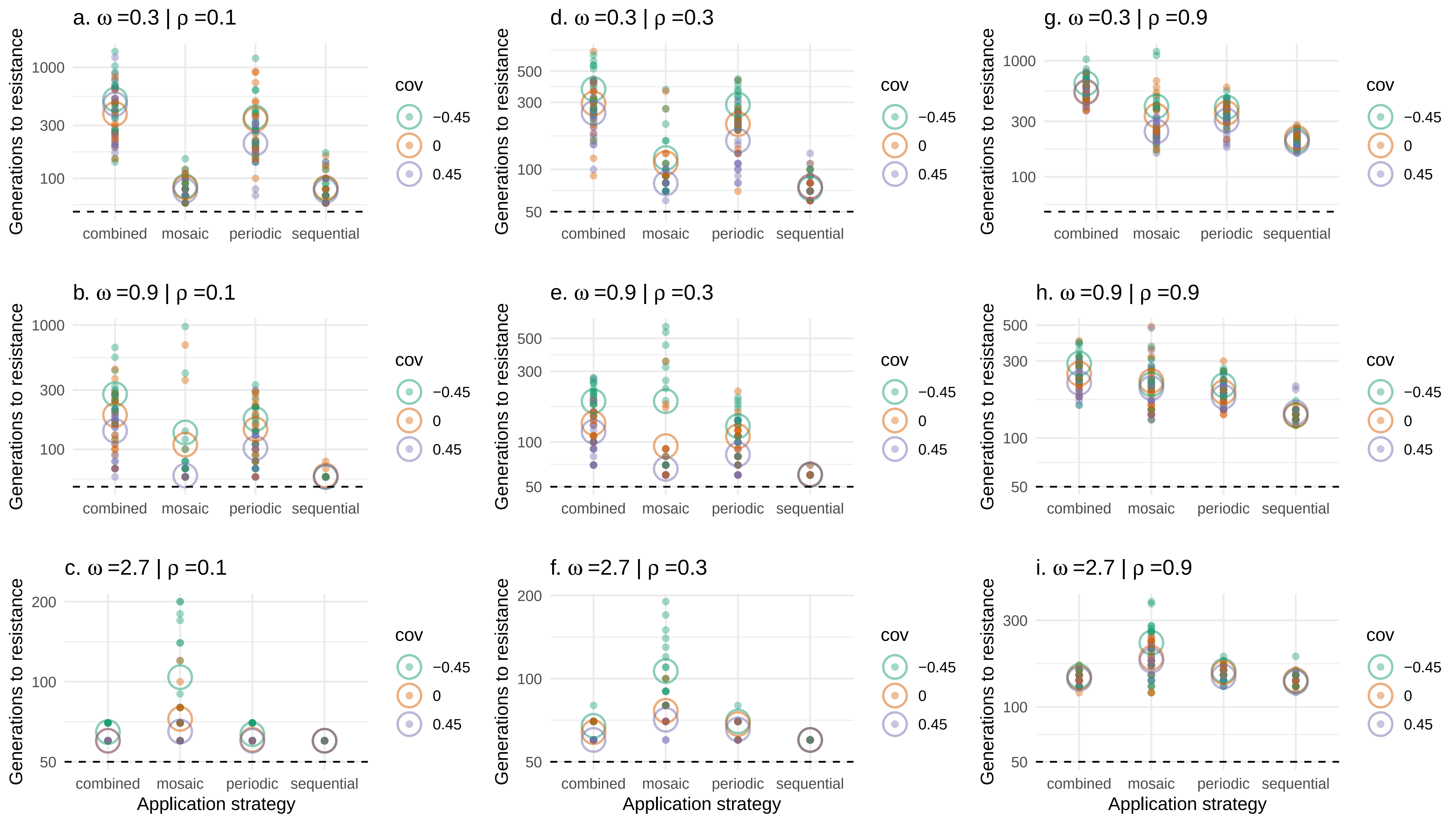

### S5

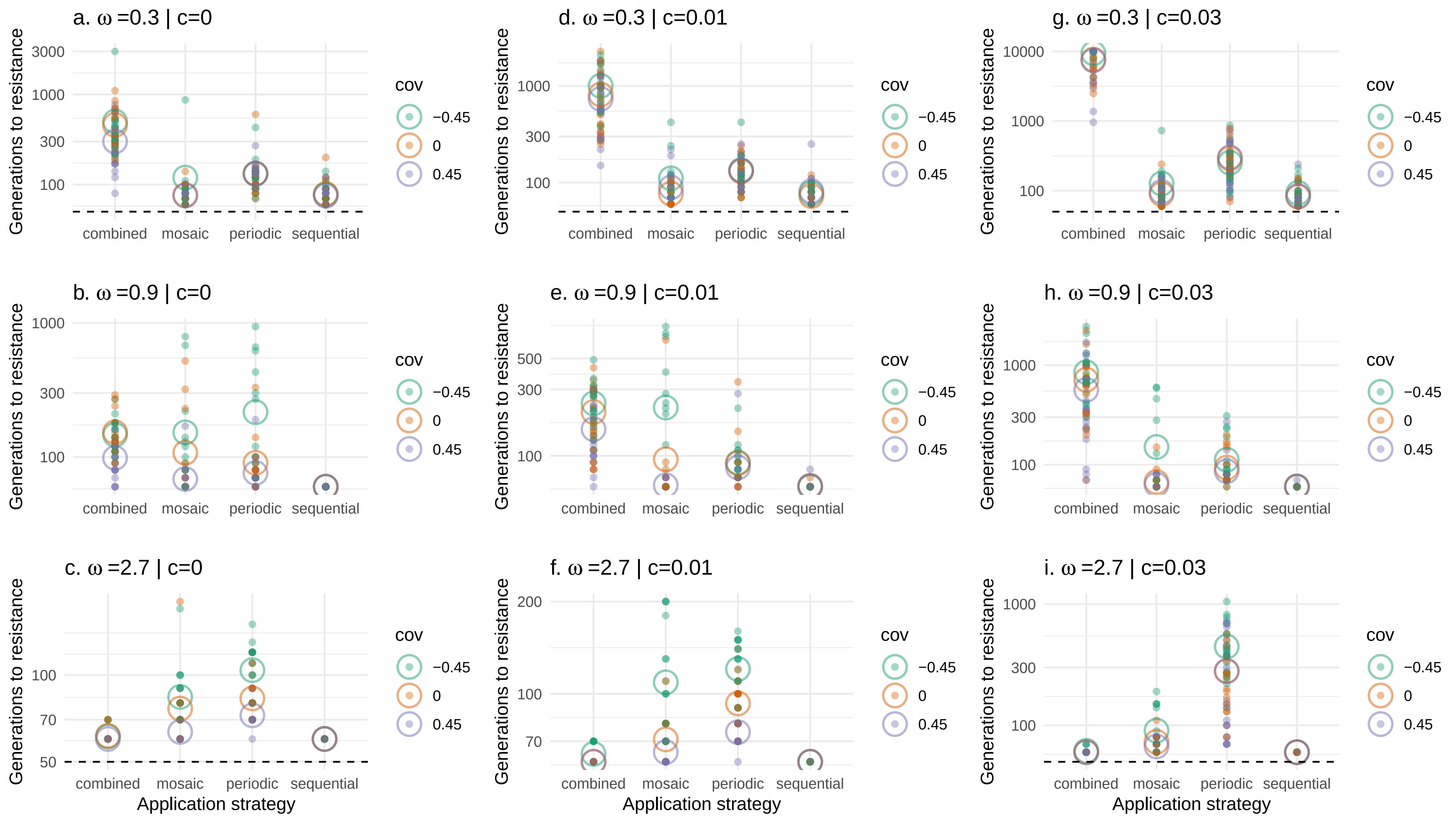
